# A cellular cross-species RNA-seq atlas captures the transcriptional dynamics of myogenesis

**DOI:** 10.64898/2025.12.03.691954

**Authors:** Fernando Valdivieso-Rivera, Anastasiia Zenina, Carlos HG Sponton, Alexander Bartelt

## Abstract

Skeletal muscle formation and regeneration is a tightly regulated process involving extensive transcriptional reprogramming as proliferating myoblasts fuse into mature myotubes. However, a comparative comprehensive analysis of the transcriptional landscape of humans and mouse myogenesis is missing. Here, we present a high-quality RNA-sequencing dataset profiling this transition in both mouse (C2C12) and human (LHCN-M2) myogenic cells. Samples were collected from proliferating myoblasts, the early differentiation phase, and from mature myocytes using identical protocols, ensuring stringent comparability. This unified dataset captures the major transcriptional shifts occurring in myoblasts, marking the onset of differentiation. Quality metrics, including PCA, read distribution, and clustering, confirmed high internal consistency across samples and species. Comparative analyses revealed shared global features of myogenesis but also distinct regulatory trajectories. Human differentiation showed early upregulation followed by suppression of metabolic and stress-related pathways, while structural and ECM-associated programs remained persistently elevated. In contrast, mouse C2C12 cells displayed early inflammatory activation and later enrichment of metabolic and contractile pathways typical of mature myotubes. Ortholog-based integration demonstrated decreasing cross-species correlation over time, indicating progressive reinforcement of species-specific differentiation programs. We additionally compared this bulk RNA-seq with tissue-derived myotubes from single-cell muscle datasets to determine which transcriptional programs are conserved in the *in vitro* models or which emerge only *in vivo*. This analysis delineates the conserved and context-specific features of myogenesis, identifying pathways that reflect culture-specific artifacts in both human and mouse muscle cells.

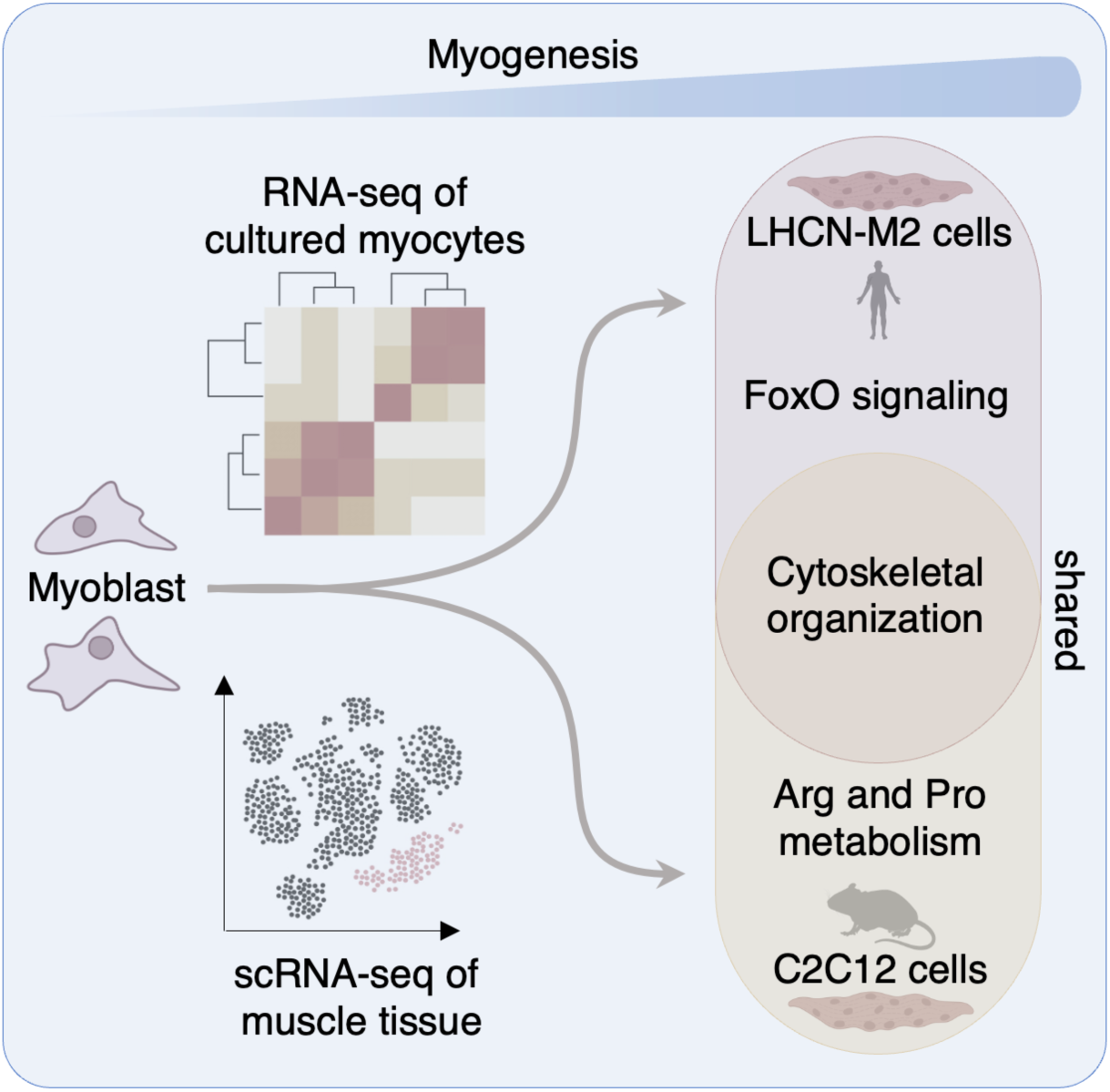

## Background & Summary

Skeletal muscle development depends on a precisely coordinated sequence of gene expression changes that regulate myoblast proliferation, lineage commitment, and fusion into multinucleated myotubes. During differentiation, transcriptional networks involving MyoD, Myogenin, Mef2, and numerous structural and metabolic genes are activated in a defined temporal order. Understanding these transcriptional dynamics is essential for elucidating the mechanisms governing muscle formation, regeneration, and disease^1,2,3^.

While mouse C2C12 cells are the most widely used model for studying myogenesis, comparable datasets from human (LHCN-M2) myogenic cells remain limited. Cross-species comparative transcriptomics can reveal evolutionarily conserved regulatory modules as well as species-specific adaptations; however, such analyses are often confounded by technical variability between sequencing batches or laboratories^4^.

### A unified cross-species dataset enables batch-reduced comparisons

To minimize batch effects, we generated a unified and directly comparable transcriptomic resource by sequencing differentiation time courses in LHCN-M2 and C2C12 under identical library preparation and analytical conditions. All libraries were prepared and sequenced by the same company using the same protocol and platform, thereby minimizing major sequencing and processing-related biases that typically hinder interspecies comparisons. Three time points were selected to represent key stages of differentiation: day 0 (myoblasts), day 1 (early differentiation, commitment phase), and day 6 (fully differentiated myotubes) for C2C12, and day 0, day 1, and day 12 for LHCN-M2, reflecting the slower differentiation kinetics of human cells (Fig. 1A). The methods used for the analysis are detailed in Figure 2B. General data quality assessments, including GC content distribution, clean read ratios, PCA clustering, and Euclidean distance among samples, as well as read mapping across genomic regions (exon, intron, and intergenic) were performed and confirmed the high quality of the data set (Fig. S1A-E). Correlation matrices visualize intra-species sample relationships for mouse and human datasets separately (Fig. S1A-E). In addition, KEGG pathway enrichment was performed using the full set of detected genes (Fig. 1C), providing an overview of the major biological processes represented in the dataset.

**Figure 1.**
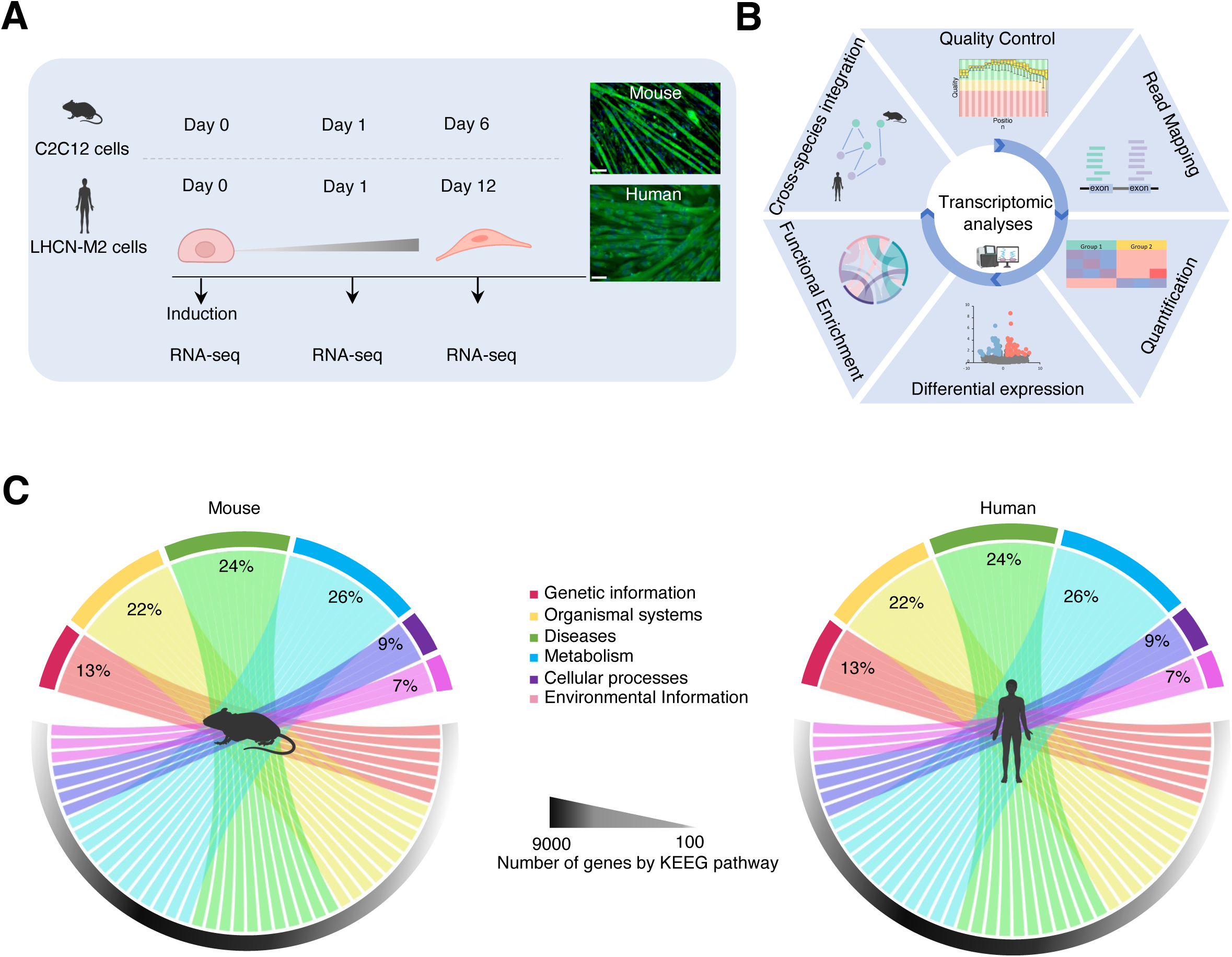
Experimental design and quality assessment of mouse and human RNA-seq datasets. (A) Schematic representation of sample collection and experimental design prior to sequencing. Scale bar = 40 μm. The cartoon was created using a Biorender. (B) Overview of the RNA-seq analytical workflow, including the bioinformatic algorithms and tools applied. The cartoon was created using a Biorender. (C) Gene Ontology (GO) chord plot illustrating the relationship between GO terms and associated genes detected in both mouse and human samples, grouped according to major KEGG pathway categories.

**Figure 2.**
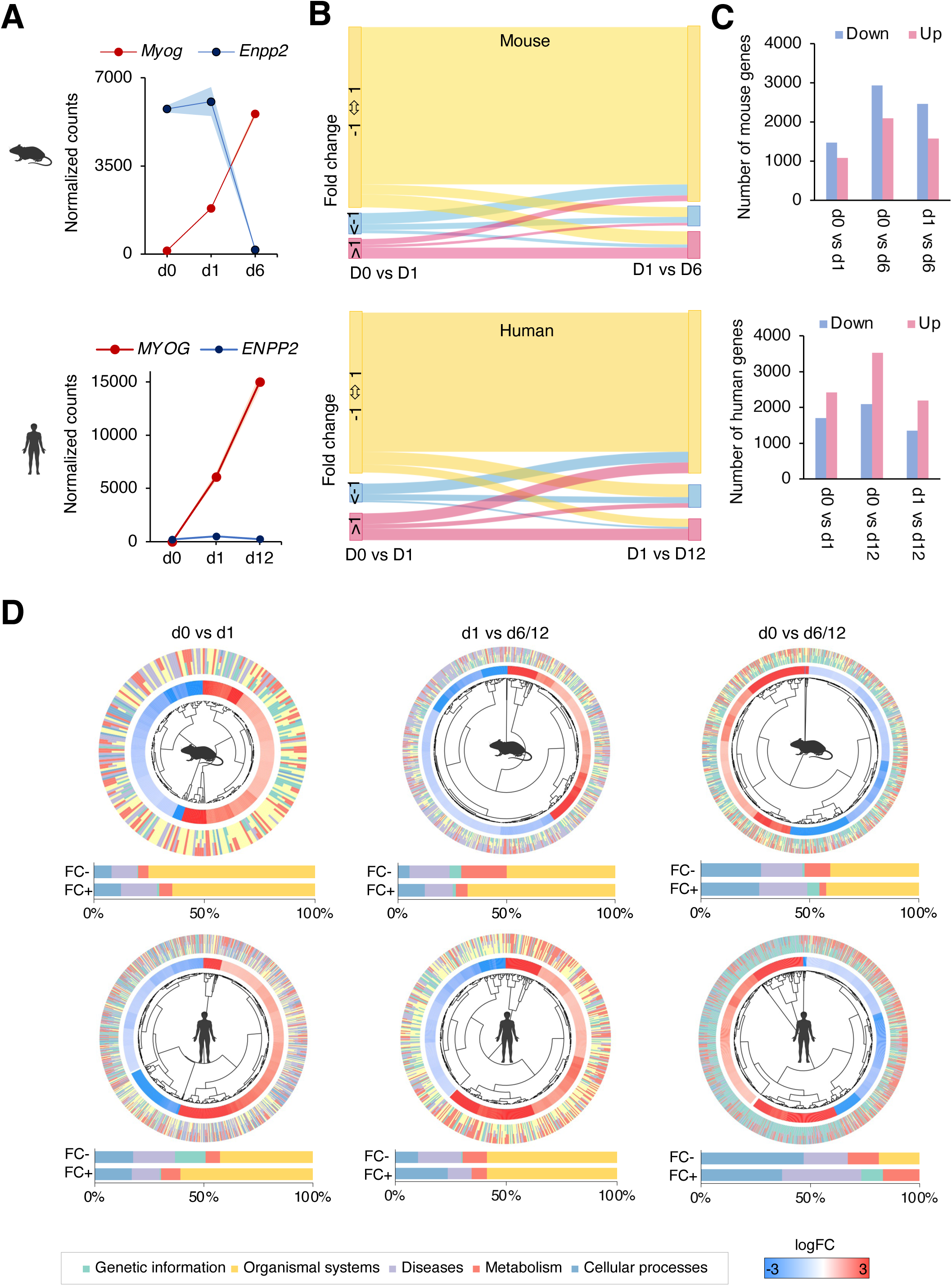
**Comparative analysis of functional enrichment and transcriptional dynamics in mouse and human myogenic differentiation.** (A) Normalized expression counts of canonical myogenic differentiation markers across different time points in mouse (top) and human (bottom) samples. (B) Sankey diagram visualizing the flow of gene expression fold changes between consecutive differentiation stages (day 0 to day 1, and day 1 to day 6), highlighting dynamic transcriptional transitions. (C) Significant differentially expressed genes identified between differentiation stages in mouse (top) and human (bottom) datasets. (D) Chord diagram showing the core genes involved in significantly enriched pathways for mouse (top) and human (bottom) samples.

### Shared and divergent temporal dynamics between mouse and human myogenesis

Sankey diagrams illustrating fold-change dynamics show that the transition from day 0 to day 1 shows the largest change among sampled intervals and bar plots summarizing the number of up- and down-regulated genes across stages, marking the onset of differentiation and major transcriptional shifts during differentiation (Fig. 2B-C). Circos plots emphasize large-scale transcriptional divergences between species, particularly between day 0 vs. day 6 in mouse and day 0 vs. day 12 in human cells (Fig. 2D). When comparing early and late stages, both species showed similar enrichment in cellular process and organismal system pathways; however, the relative contribution of these categories varied across time points. In early stages (day 0 vs. day 1), cellular process-related genes represented a larger fraction of the differentially expressed set, while in later stages, organismal system genes became more prominent. Interestingly, this pattern was inverted in humans when comparing day 1 vs. day 12, suggesting distinct temporal coordination of differentiation programs between species (Fig. 2D). Together, these results indicate that organismal process-related genes are consistently the most dynamically regulated across differentiation, except for at day 12 in humans, and that interspecies differences may arise from distinct cell type-specific regulatory dynamics.

### Human myogenesis undergoes sequential loss of signaling programs and consolidation of structural networks

To further explore these differences, we compared pathway transitions across stages in human myogenic differentiation. Several pathways upregulated between day 0 and day 1, including cholesterol metabolism, ferroptosis, and steroid biosynthesis, became downregulated between day 1 and day 12, reflecting metabolic rewiring during maturation. In contrast, classical muscle-related pathways such as cytoskeletal organization, calcium signaling, and ECM-receptor interaction remained persistently upregulated (Fig. 3A). At the category level, the percentage of downregulated genes in day 0 vs. day 12 was threefold higher than in day 0 vs. day 1, while the fraction of upregulated genes remained stable, suggesting that specific signaling programs are progressively lost during differentiation, whereas structural and transport-related genes remain active (Fig. 3A).

**Figure 3.**
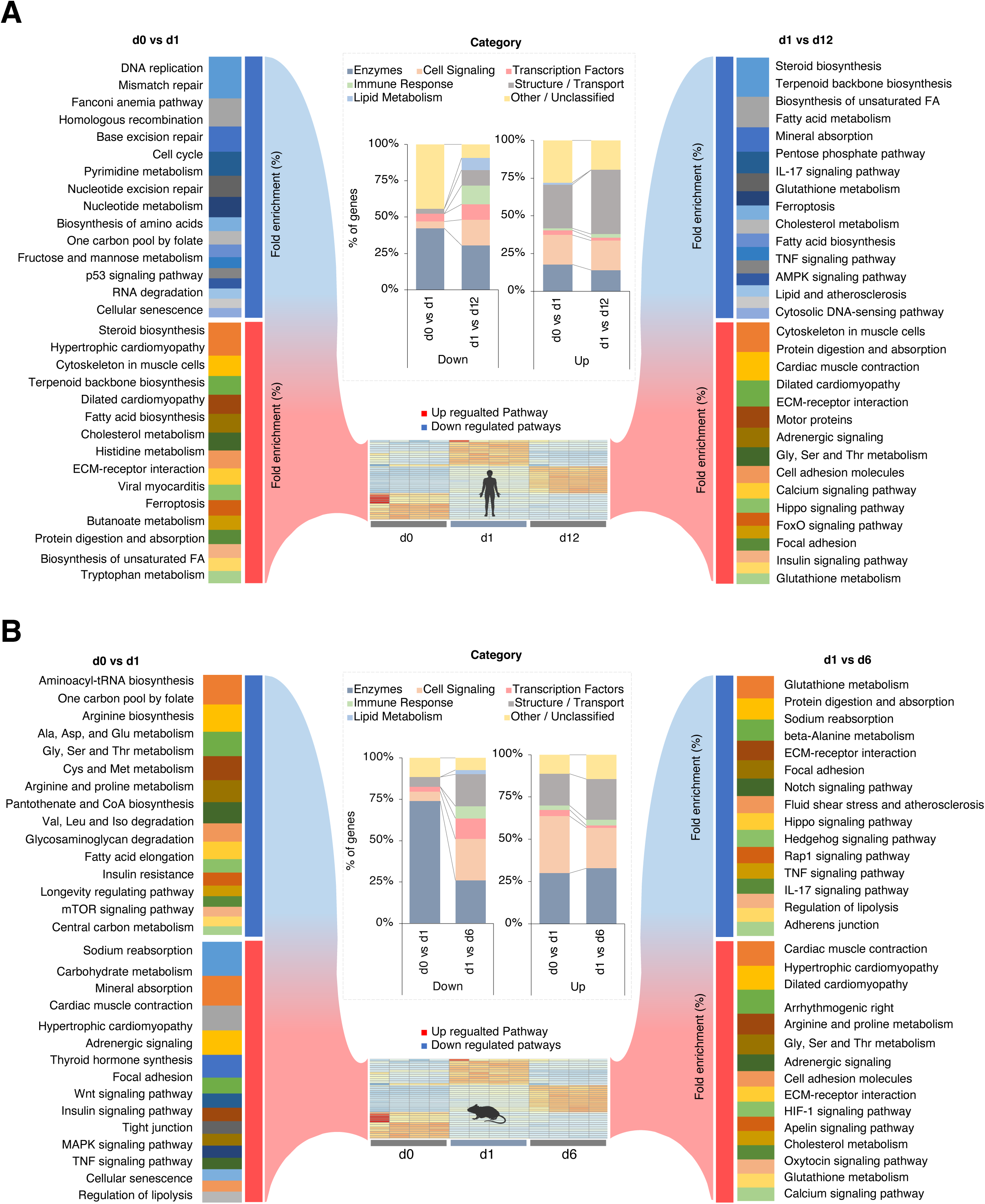
**Enriched pathways and functional gene categories in human and mouse.** (A) Enriched pathways in human genes, showing fold enrichment for up- and down-regulated pathways across two time points during differentiation. Functional categories of the genes are shown in the center of the figure as percentages. (B) Enriched pathways in mouse genes, showing fold enrichment for up- and down-regulated pathways across two time points during differentiation. Functional categories of the genes are shown in the center of the figure as percentages.

### Mouse myogenesis shows early inflammatory activation and late metabolic–contractile maturation

In mouse C2C12 cells, the most significantly enriched pathways were related to amino acid metabolism. In early stages, inflammatory pathways such as TNF-α and IL-17 signaling were upregulated, but downregulated in late stages, indicating their role during early differentiation. Conversely, pathways typical of mature muscle, such as calcium signaling and cardiac muscle contraction, became more enriched at later stages (Fig. 3B). Enzymatic genes were the predominant downregulated category during early differentiation, but this percentage decreased in later stages, whereas upregulated enzymes remained stable, consistent with their essential roles in terminal differentiation. Importantly, the proportion of genes associated with structural components remained constant, in contrast to the pattern observed in human cells (Fig. 3B). Several pathways were conserved between human and mouse myogenesis, including ECM-receptor interaction, glutathione metabolism, glycine/serine/threonine metabolism, and adrenergic signaling, supporting the presence of a core differentiation program shared across species. Notably, 10-42 % of the differentially expressed genes lacked clear functional annotation, highlighting the presence of potentially novel regulators of myogenesis.

### Ortholog integration reveals increasing species-specific divergence over time

An integrative cross-species analysis of orthologous genes log_2_(TPM+1) revealed a positive overall correlation between mouse and human gene expression, although the correlation coefficient progressively declined with differentiation time (Fig. 4A). This trend suggests that as differentiation proceeds, species-specific genes increasingly shape the transcriptional landscape. Clustering analyses identified groups of genes with high and low interspecies concordance (Fig. 4B-C), revealing divergent regulatory modules that underlie unique aspects of mouse and human myogenesis. Correlation heatmaps (Fig. 5A-C) further uncovered clusters of co-expressed and inversely correlated genes, indicating both conserved and divergent transcriptional programs. Pathway analysis of these clusters showed that highly correlated genes at day 0 shared FOXO signaling, while at day 1, overlapping pathways included cell cycle regulation, glycolipid metabolism, and cytoskeletal organization. By day 6/12, several pathways appeared inversely regulated up in mouse but down in human cells, and vice versa, indicating that distinct transcriptional strategies can achieve similar differentiation outcomes.

**Figure 4.**
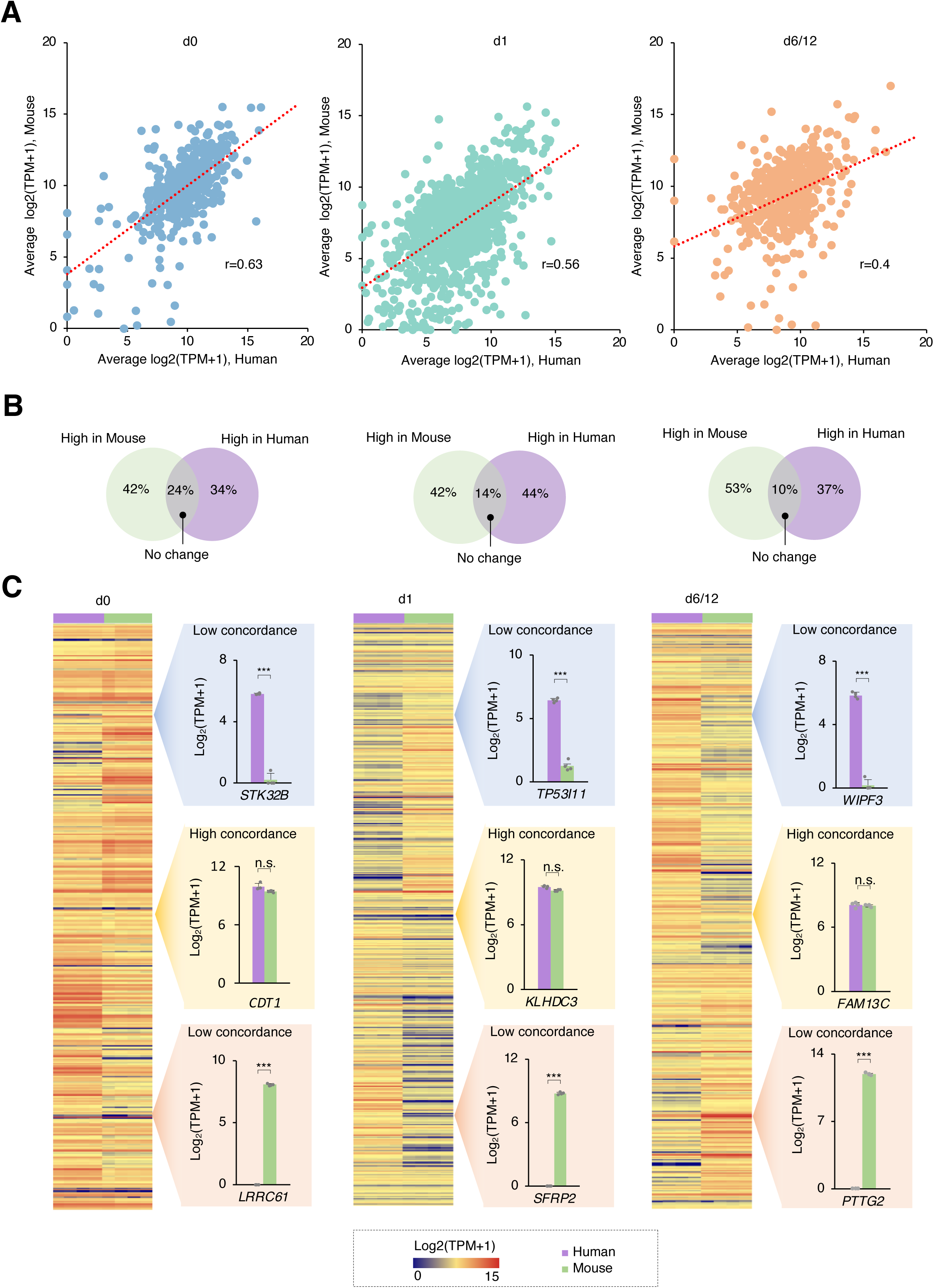
**Cross-species integrative transcriptomic analysis reveals conserved and divergent gene expression patterns during differentiation.** (A) Correlation plots of average log₂(TPM + 1) values for orthologous genes between mouse and human samples at day 0, day 1, and day 6. The regression trend lines indicate an overall positive correlation, with a gradual decrease in correlation strength as differentiation progresses. Pearson correlation coefficients between human and mouse gene expression for the three groups are shown lower right corner of each panel. (B) Venn-style diagram showing the percentage of genes assigned to each expression category when comparing human and mouse myogenesis. (C) Heatmaps showing the expression of orthologous genes (log₂(TPM + 1)) in human and mouse samples at day 0, day 1, and day 6. Genes were grouped into three expression concordance categories:low concordance (high in human, low in mouse), low concordance (high in mouse, low in human), and high concordance (similar expression levels in both species).

**Figure 5.**
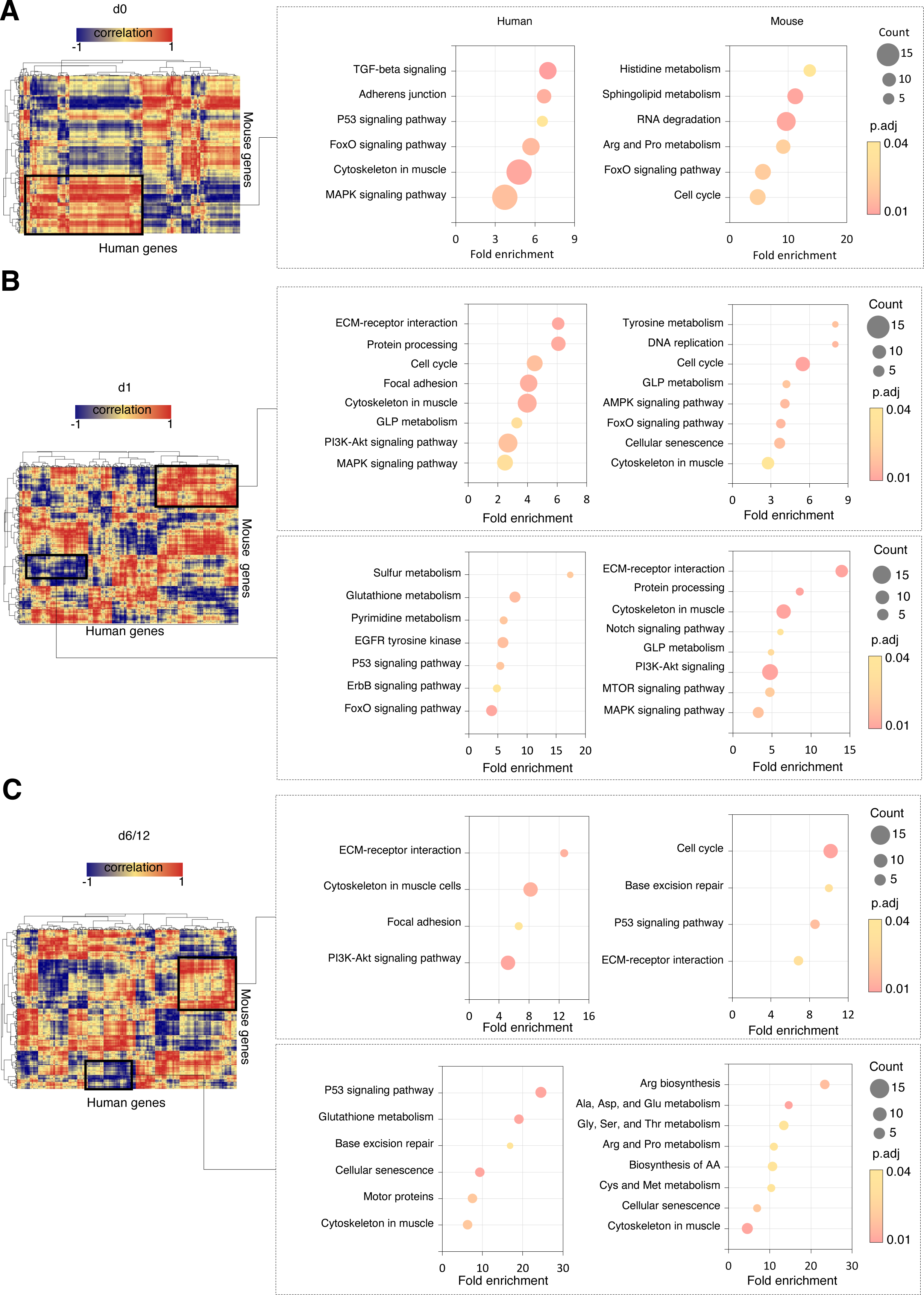
**Correlation-based clustering highlights conserved and species-specific gene modules.** (A) Left panel: heatmap of gene–gene correlations calculated from log₂(TPM+1) values in Day 0, with clusters highlighted in red indicating high positive correlation and in blue indicating low or negative correlation. Right panel: GO pathway enrichment analysis for genes within each cluster, shown separately for human and mouse, highlighting the clusters indicated in the heatmap. (B) Left panel: heatmap of gene–gene correlations calculated from log₂(TPM+1) values in Day 1, with clusters highlighted in red indicating high positive correlation and in blue indicating low or negative correlation. Right panel: GO pathway enrichment analysis for genes within each cluster, shown separately for human and mouse, highlighting the clusters indicated in the heatmap. (C) Left panel: heatmap of gene–gene correlations calculated from log₂(TPM+1) values in Day 6/12, with clusters highlighted in red indicating high positive correlation and in blue indicating low or negative correlation. Right panel: GO pathway enrichment analysis for genes within each cluster, shown separately for human and mouse, highlighting the clusters indicated in the heatmap.

### Parallel analyses in mice and humans reveal divergent structural, metabolic, and regulatory programs between tissue- and culture-derived myotubes

To compare intrinsic myotube transcriptional programs with those present in native skeletal muscle, we first selected and isolated differentiated myotubes from a high-resolution single-cell RNA-seq muscle dataset^6^ and normalized their expression profiles to enable the direct comparison to our bulk RNA-seq measurements in C2C12 cells (Fig. 6A). To evaluate the magnitude of expression across platforms, all genes were ranked within each dataset and stratified into quartiles. Genes above the 75^th^ percentile were classified as highly expressed, whereas those below the 25^th^ percentile were considered lowly expressed. This framework allowed us to identify genes that were consistently high in both systems, as well as those showing preferential expression in either *in vitro* myotubes or *in vivo* myotubes.

**Figure 6.**
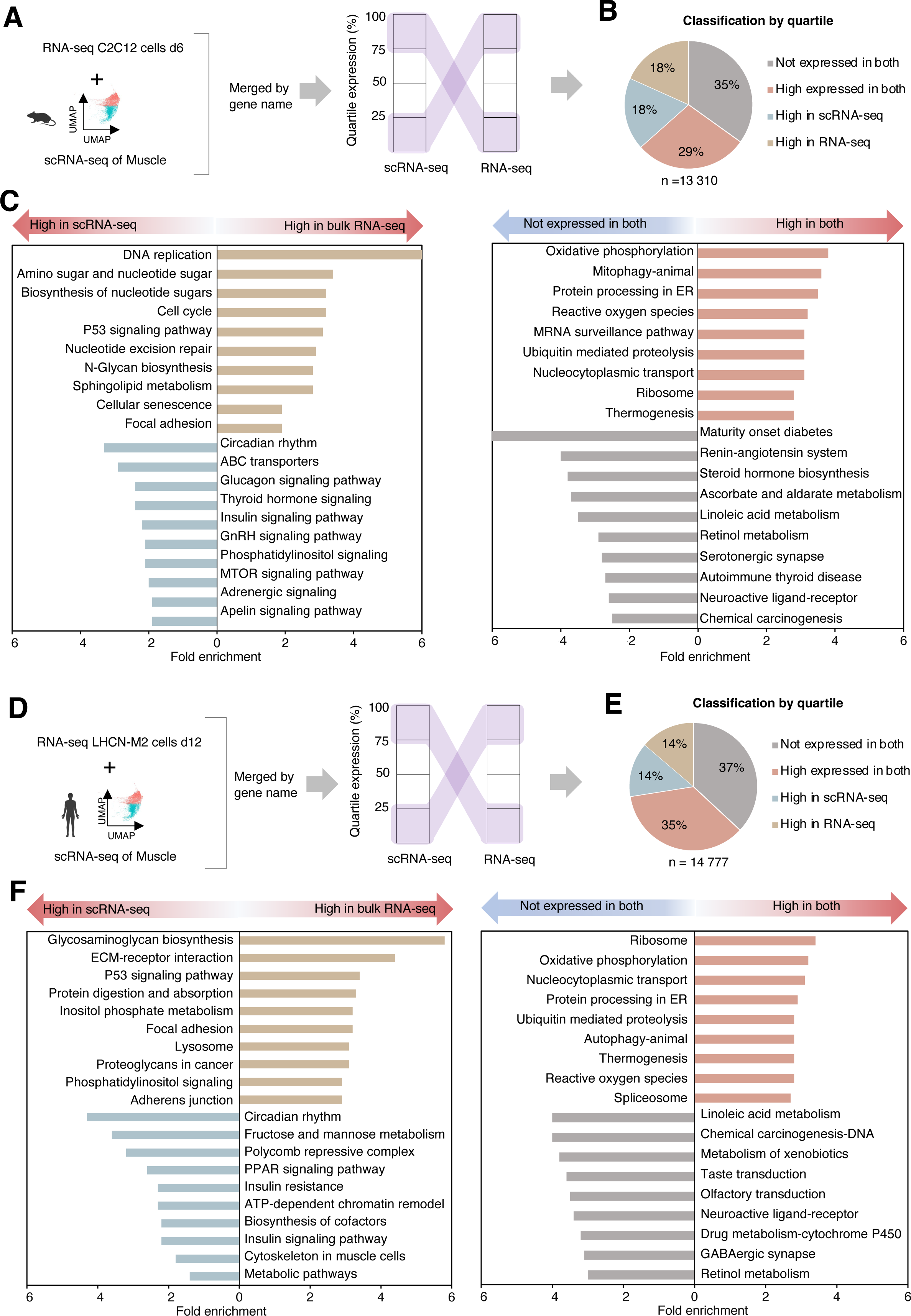
**Integration of bulk and single-cell RNA-seq datasets in mouse and human myotubes.** (A) Schematic overview of the workflow used to integrate bulk RNA-seq and single-cell RNA-seq datasets from mouse samples. (B) Percentage-based comparison of the curated gene set across datasets, highlighting shared and dataset-specific expression. (C) Enriched GO metabolic pathways distinguishing bulk RNA-seq and single-cell myotube profiles. (D) Schematic overview of the workflow used to integrate bulk RNA-seq and single-cell RNA-seq datasets from human samples. (E) Percentage-based comparison of the curated gene set across datasets, highlighting shared and dataset-specific expression. (F) Enriched GO metabolic pathways distinguishing bulk RNA-seq and single-cell myotube profiles.

Across the 13,310 curated genes examined, 29 % genes were highly expressed in both datasets, indicating a conserved myotube transcriptional core shared between cultured and tissue-derived myocytes. In contrast, two distinct gene sets of similar size (18% of genes in each group) showed asymmetric expression: One set was selectively enriched in bulk C2C12 myotubes, whereas the other was preferentially enriched in the single-cell muscle-derived myotubes (Fig. 6B. This separation reflects fundamental biological differences between proliferative, *in vitro* differentiated myotubes and mature myofibers embedded in a physiological tissue environment.

Pathway analysis of the C2C12-enriched genes revealed strong representation of proliferative and biosynthetic programs, including DNA replication, cell-cycle progression, nucleotide-sugar and N-glycan biosynthesis, nucleotide-excision repair, sphingolipid metabolism, cellular senescence, and focal-adhesion remodeling. These pathways are characteristic of immortalized myogenic cultures undergoing sustained growth, membrane construction, and high biosynthetic activity. In contrast, genes enriched in single-cell derived myotubes mapped to metabolic and endocrine regulatory circuits, including circadian rhythm, insulin, glucagon and thyroid hormone signaling, the PI3K/MTOR axis, adrenergic and apelin pathways, ABC transporters and GnRH signaling (Fig. 6C). These signatures capture the hormonally integrated and metabolically coordinated state of mature *in vivo* muscle fibers, in which systemic signals and rhythmic metabolic inputs shape transcriptional activity.

We also performed the same analysis using single-cell RNA-seq^7^ data from muscle humans (Fig. 6D). Of 14,777 curated genes examined, 35 % genes were highly expressed in both datasets, indicating a conserved myotube transcriptional core shared between cultured and tissue-derived muscle (Fig. 6E). Bulk RNA-seq was enriched for structural and homeostatic pathways, including extracellular-matrix organization (glycosaminoglycan biosynthesis, ECM-receptor interaction, focal adhesion), adhesion processes (adherents’ junction), lysosomal activity, p53 signaling, and phosphatidylinositol metabolism. In contrast, human single-cell data highlighted dynamic regulatory and metabolic programs, such as circadian rhythm, insulin and PPAR signaling, chromatin remodeling, carbohydrate metabolism, and broader metabolic pathways (Fig. 6F). Overall, in both species, bulk measurements emphasize structural, biosynthetic, or stress-related programs, whereas single-cell profiles more clearly capture endocrine, metabolic, and regulatory signaling, consistent with higher sensitivity to active transcriptional states.

### Significant statement

By generating matched bulk RNA-seq datasets from human muscle LHCN-M2 and C2C12 myotubes produced within the same technical and analytical environment, this study minimizes batch-related confounders and enables a robust cross-platform comparison. When contrasted against single-cell resolved myotube profiles obtained from intact muscle *in vivo*, our analysis reveals that several pathways strongly enriched in C2C12 cells, such as replication-associated and amino-sugar biosynthetic programs, are largely absent in mature muscle *in vivo*. This framework identifies which transcriptional features likely reflect conserved myogenic biology and which could arise as culture-specific artifacts. The datasets provided here offer a practical reference for selecting genes and pathways a with higher likelihood of *in vivo* translatability, ultimately guiding more rigorous experimental design in muscle and metabolic research.

Importantly, this cross-platform comparison also provides a cautionary note for the interpretation of replication analyses performed exclusively in cell culture. Several pathways that appear robustly activated in C2C12 myotubes, such as DNA replication, nucleotide-sugar biosynthesis, and focal-adhesion remodeling, are largely decreased from the transcriptional landscape of mature myofibers *in vivo*. Conversely, key regulatory circuits that dominate the native muscle environment-hormone-responsive metabolic pathways, circadian regulation and nutrient-sensing signaling are not prominently found under standard *in vitro* conditions. These discrepancies indicate that replication of molecular effects in C2C12 cells does not necessarily equate to physiological relevance and highlight the importance of validating *in vitro* findings from these cell lines rigorously. By integrating bulk and single-cell muscle transcriptomes, our analysis delineates which aspects of myogenic biology are conserved across contexts and which emerge only within the systemic, multicellular environment of intact muscle.

To more accurately replicate the endocrine and metabolic cues found *in vivo*, future research should investigate the deliberate addition of systemic signaling factors to differentiation media, especially in the final stages of myogenesis. Our analysis indicates that wider integration of hormonal, nutrient-sensing, and circadian regulators could significantly improve the physiological fidelity of *in vitro* cultured myotubes, even though some compounds are already used to mimic specific metabolic pathways. By incorporating these systemic cues, it may be possible to close the remaining gap between tissue-derived muscle myotubes and cultured myotubes, allowing for more precise modeling of the biology of human and mouse muscle function in cultured cells.

## Methods

### >Human skeletal muscle cell culture and differentiation

Human skeletal myoblasts (LHCN-M2; CkHT-040-231-2, Everycite) were maintained in growth medium consisting of MyoUp ready-to-use medium (C-28080, PromoCell) or a custom mixture of DMEM (Gibco, Cat# 61965-026) and M199 (Gibco, Cat# 31150-022) at a 4:1 ratio, supplemented with 15% fetal bovine serum (PAN Biotech, Cat# P30-3031), 20 mM HEPES (Sigma-Aldrich, Cat# H0887), 3 μg/ml zinc sulfate (Sigma-Aldrich, Cat# Z0251), 1.4 μg/ml vitamin B12 (Sigma-Aldrich, Cat# V2876), 0.055 μg/ml dexamethasone (Sigma-Aldrich, Cat# D4902), 2.5 ng/ml hepatocyte growth factor (HGF; Merck Millipore, Cat# GF116), and 5 ng/ml basic fibroblast growth factor (bFGF; Enantis, Cat# FGF-STAB). Cells were cultured on collagen-coated dishes (0.1% gelatin solution, Sigma-Aldrich) at 37 °C in a humidified atmosphere containing 5% CO2, and passaged at 70–80% confluence using Trypsin-EDTA (0.25%) (Gibco). For differentiation, cells were seeded at 90% confluence in culture plates pre-coated with 0.1% gelatin. Differentiation medium consisted of DMEM/M199 (4:1) supplemented with 2% horse serum. To prepare the medium, 8 ml DMEM (Gibco, 10566-016) and 2 ml M199 (Gibco, 31150-022) were combined in a sterile 15-ml tube and mixed thoroughly. After discarding 200 µl of the mixture to maintain the final volume ratio, 200 µl horse serum (Sigma-Aldrich, H1270) was added and gently mixed. The medium was switched every 48 hours for up to 12 days.

### >C2C12 myoblast culture and differentiation

Cells were differentiated as previously described^7^. Mouse myoblasts (C2C12) were maintained in growth medium composed of Dulbecco’s Modified Eagle Medium (DMEM; Gibco) supplemented with 10% fetal bovine serum and 1% penicillin-streptomycin (Gibco). Cells were cultured on tissue-culture dishes at 37 °C in a humidified atmosphere containing 5% CO2, and passaged every 2-3 days before reaching 80% confluence to preserve their myogenic potential. For differentiation, cells were seeded at 90% confluence and switched to differentiation medium containing DMEM supplemented with 2% horse serum (Gibco). Medium was replaced every 48 h, and differentiated myotubes were harvested at the indicated time points.

### >RNA extraction and sequencing

Total RNA was extracted using TRIzol reagent (Thermo Fisher Scientific). The supernatant was transferred to NucleoSpin RNA purification columns for subsequent extraction and on-column DNase treatment. RNA integrity was assessed with an Agilent 2100 Bioanalyzer, and samples with RNA integrity number (RIN) ≥ 8.0 were used. Library preparation and sequencing for all samples were performed by the same sequencing company using a standardized protocol. Sequencing was performed on the Illumina Novaseq platform (strategy PE150) by Novogene Co., Ltd.

### >Data processing and quality control

Raw reads were first evaluated for quality using FastQC^8^. Trimmomatic^9^ was additionally used to remove adapter sequences, trim low-quality bases, and discard reads falling below quality thresholds. Clean reads were aligned to the mouse (mm10) or human (GRCh38) reference genomes using HISAT2^10^ with default parameters. Aligned reads were assembled into transcripts with StringTie^11^, and gene-level quantification was performed using featureCounts^12^. Quality control metrics (GC content, mapping efficiency, and genomic feature distribution) were summarized across samples. PCA and Euclidean distance analyses were then performed using normalized TPM values to evaluate sample clustering and overall dataset structure.

### >Differential expression and functional analysis

Differential expression analysis was performed with DESeq2^13^ using adjusted p-value < 0.05 and |log2FC| ≥ 1. Gene Ontology (GO)^14^ and KEGG^15^ pathway enrichment analyses were carried out using clusterProfiler. Sankey diagrams and bar plots were generated with the ggalluvial and ggplot2 packages. Circos plots were constructed using the R package circlize to visualize species-specific transcriptional differences. Orthologous genes were identified through Ensembl BioMart, and integrated analyses were performed using log_2_(TPM + 1) matrices.

### >Single-cell RNA-seq data reanalysis

Publicly available single-cell RNA-seq count matrices and metadata from the original study were imported into Seurat (v5) in R. Myotubes were isolated strictly according to the cell-type annotation provided by the original authors in the metadata; that is, we subset cells labeled as “myotube” without reassigning cell identities. For cross-platform comparisons, expression values were aggregated using AverageExpression and transformed to log_2_(CPM + 1). Genes were ranked within each dataset and binned into quartiles to classify high- and low-expression groups. When integrating species, only one-to-one orthologs were retained. To enable a scale-independent comparison of relative expression strength, genes within each dataset were ranked by their normalized expression and stratified into four quartiles: 0-25th percentile (low), 25-50th (moderate-low), 50-75th (moderate-high), and 75-100th percentile (high expression). This quartile-based classification provides a robust, distribution-aware framework that avoids biases introduced by absolute expression thresholds, allowing us to identify genes consistently expressed at similar relative levels across datasets and to distinguish those preferentially enriched in either bulk-differentiated or tissue-derived myotubes.

### >Data Records

Raw and processed RNA-seq data for both mouse and human samples have been deposited in the Gene Expression Omnibus (GEO) under accession number [to be assigned]. Deposited files include: Raw FASTQ files for each sample (C2C12: day 0, 1, 6; LHCN-M2: day 0, 1, 12). Metadata file including species, day, replicate number, and codes.

### >Technical Validation

Sequencing depth averaged million reads per sample with >97% high-quality bases. GC content distributions were uniform across samples (Fig. S1). PCA and Euclidean distance analyses confirmed clear separation by time point and high reproducibility among biological replicates (Fig. S1). Differential expression revealed that the transition from day 0 to day 1 accounted for the most pronounced transcriptional reorganization in both species, consistent with the initiation of differentiation. KEGG analysis showed enrichment of pathways related to muscle contraction, mitochondrial metabolism, and extracellular matrix organization. Circos visualization highlighted divergent regulation in metabolic and cell adhesion pathways between mouse and human datasets. Cross-species ortholog mapping revealed clusters of conserved myogenic gene expression. Overall, the dataset demonstrates high technical consistency and strong biological signal.

### >Usage Notes

The deposited data can be directly used for downstream analyses, including alternative splicing detection, isoform quantification, or cross-species network inference. All raw FASTQ files can be reprocessed using alternative pipelines if desired. For comparative purposes, we used log_2_(TPM + 1) or Log_2_(CPM + 1) transformation and Z-score normalization before analyses. Scripts available in the GitHub repository include automated pipelines for quality control, normalization, differential expression, KEGG enrichment, and figure generation (PCA, Sankey, circos, and heatmaps).

## Code Availability

All code and figure-generation scripts are available in the GitHub repository [link], including R and Python scripts for data processing, normalization, and visualization. The repository also contains example workflows to reproduce the analyses and figures presented in this paper.

## Lead contact

Further information and requests for materials should be directed to and will be fulfilled by the lead contact Alexander Bartelt.

## Material availability

Materials generated in this study are available upon request from the lead contact.

## Author contributions

Conceptualization, F. V. R., A.Z., and A.B; investigation, F.V.R and A.Z; writing original draft, F.V.R; review and editing A.Z., C.H.S and A.B.; funding acquisition, C.H.S. and A.B.

## Acknowledgments

We thank Marlene Stumbaum for excellent technical assistance and all members of the Bartelt Lab for the enjoyable atmosphere. This work used resources of the Centro Nacional de Processamento de Alto Desempenho em São Paulo (CENAPAD-SP). This work was supported by the São Paulo Research Foundation FAPESP (2024/13043-4, 2019/15025-5) to F.V.R and C.H.S., and by the Else Kröner Fresenius Foundation (EKFS), by the German Center for Cardiovascular Research (DZHK, 81X2600423) and by the European Research Council (ERC) Starting Grant PROTEOFIT 852742 to A.B.

---

**Figure S1.**
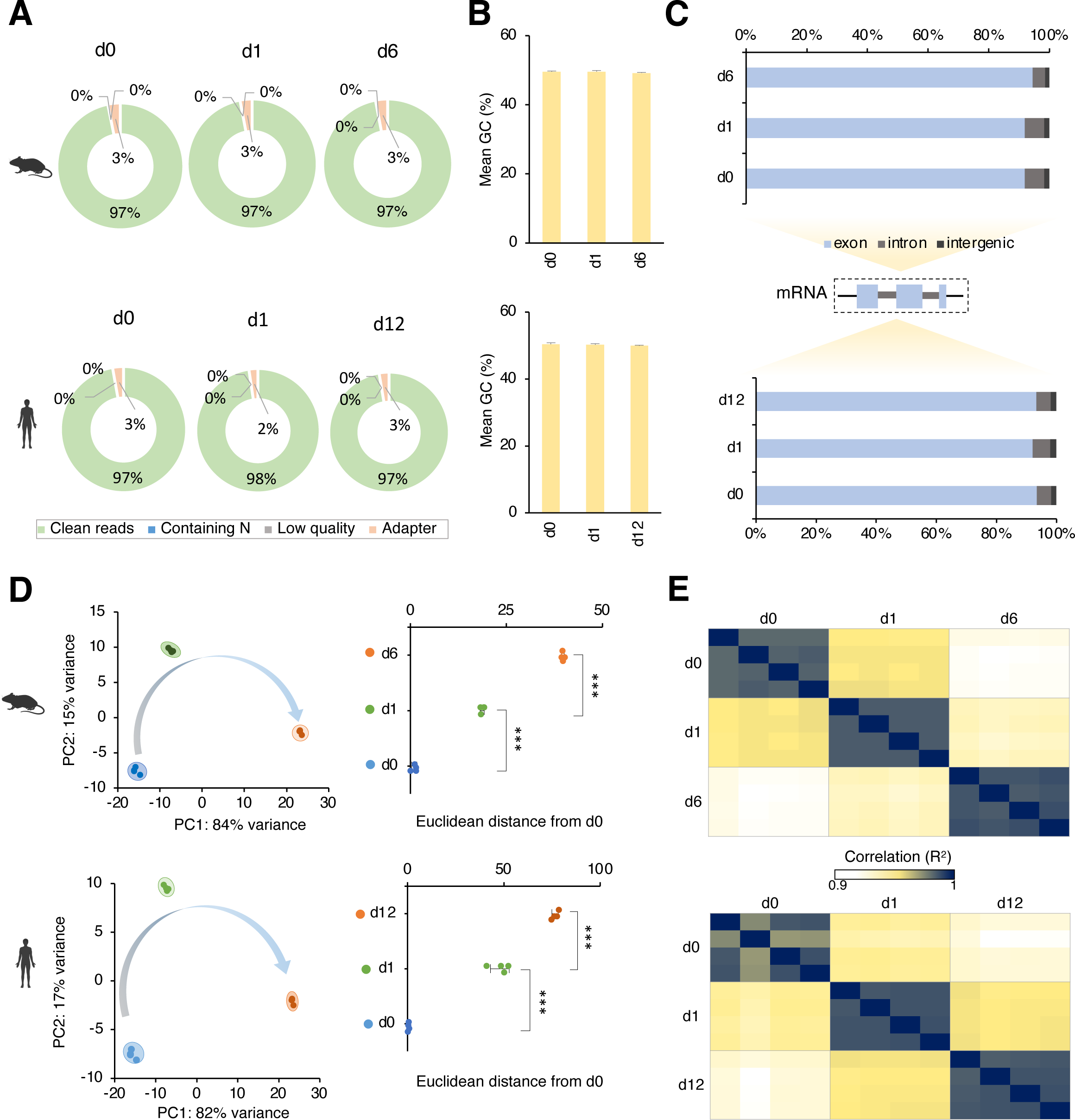
**Quality assessment of mouse and human RNA-seq datasets.** (A) Average percentage of clean reads obtained per differentiation day for mouse (top) and human (bottom) samples. (B) GC content percentage across differentiation stages for mouse (top) and human (bottom) samples. (C) Distribution of mapped reads across exonic, intronic, and intergenic regions for mouse (top) and human (bottom) samples. (D) Principal component analysis (PCA) illustrating the separation of samples across differentiation days. The Euclidean distance plot on the left represents sample divergence relative to day 0. (E) Pearson correlation matrices displaying transcriptomic similarity among samples, with mouse (top) and human (bottom) datasets. For B and D, data are presented as the mean±SEM. ***P < 0.001.

